# Optimizing automated sleep stage scoring of 5-second mini-epochs: a transfer learning study

**DOI:** 10.1101/2025.06.28.661238

**Authors:** Louise Frøstrup Follin, Julie Anja Engelhard Christensen, Janita Vevelstad, Hilde T. Juvodden, Rannveig Viste, Berit Hjelde Hansen, Mathias Perslev, Tobias Kaufmann, Al-exander Neergaard Zahid, Stine Knudsen-Heier

**Author notes:** Corresponding author: Louise Frøstrup Follin, Norwegian Centre of Expertise for Neurodevelopmental Disorders and Hypersomnias (NevSom), Oslo University Hospital, Box 4956 Nydalen, 0424 Oslo, Norway. These authors have contributed equally (Al-exander Neergaard Zahid, Stine Knudsen-Heier).

## Abstract

**Study objective:** Conventional sleep staging relies on 30-second epochs, potentially concealing transient sleep stage intrusion and reducing precision. Building on our previous study of mini-epochs, we investigated whether U-Sleep, an existing automatic deep learning-based sleep staging model with high performance in epochs, could be optimized to similar performance level in 5-second mini-epoch scoring, thereby enabling more detailed sleep characterization.

**Methods:** We created a dataset of 48,000 human-scored 5-second mini-epochs from 100 PSGs. We compared mini-epochs to human-scored epochs before U-Sleep was optimized using transfer learning and evaluated on a test set. Model performance was assessed using F1-scores, confusion matrices, stage distributions and transition rates comparing scorings of the original U-Sleep before, and the optimized U-Sleep after transfer learning to human-scored mini-epochs.

**Results:** Compared to human-scored epochs, human-scored mini-epochs captured significantly more transitions (1.70/minute vs. 0.21/minute, p<0.001), and significantly more wake (8.4% versus 5.4%), N1 (7.2% versus 5.4%), and N2 (51.8% versus 40.9%), less N3 (15.4% versus 25.2%) and REM sleep (16.7% versus 23.0%) (all p<0.001). Optimizing U-Sleep improved its performance significantly from F1=0.74 to F1=0.81 (p<0.05) and gave increased transition rates in the test set (original U-Sleep: 1.06/minute, optimized U-Sleep: 1.34/minute, human-scored mini-epochs: 1.70/minute). Stage distributions did not differ between optimized U-Sleep’s scorings and human-scored mini-epochs.

**Conclusions:** After optimization, U-Sleep performance in mini-epochs matched the high performance levels previously reported in both human and automated 30-second epoch scoring. This demonstrates the feasibility of precise, automated high resolution sleep staging. Future work should include external validation and application to full-night recordings.

**Statement of significance:** Conventional 30-second epochs limit temporal resolution in sleep staging and may conceal transient intrusions of wake or sleep stages. However, no validated methods are available for highresolution scoring. In this study, we trained and validated the state-of-the-art deep learning model U-Sleep for accurate automatic 5-second mini-epoch scoring using a large dataset of human-scored mini-epochs. The optimized model achieved a high performance, matching levels from previously reported automatic and human epoch scoring. Compared to epoch scoring, mini-epochs captured significantly more stage transitions, supporting their ability to uncover sleep dynamics that are otherwise lost. Our findings show the potential of high-resolution sleep staging for more detailed characterization of sleep architecture and demonstrate the feasibility of precise, automatic mini-epoch scoring.

## Introduction

Accurate sleep staging is fundamental for the assessment of sleep disorders and the understanding of sleep physiology. Traditionally, sleep stages are scored manually in 30-second epochs based on visual evaluation of polysomnography (PSG) recordings. Early sleep staging relied on paper recordings, where each paper sheet could include 30 seconds, establishing the 30-second epoch length (1). The first sleep stage scoring manual was defined by Rechtschaffen and Kales (R&K) in 1968 (2). The American Academy of Sleep Medicine (AASM) updated the scoring guidelines in 2007 (3), refining staging criteria while maintaining the 30-second epoch length for simplicity (4,5).

However, the fixed 30-second epoch length does not reflect the continuous and dynamic nature of sleep, and characteristics from multiple sleep stages can appear within a single 30-second epoch. The AASM guideline addresses this issue by instructing scorers to classify an epoch based on the predominant sleep stage (3). While this approach potentially simplifies scoring, it may also contribute to interrater variability when scorers for example need to decide which of the potentially several present sleep stages fills the majority of the epoch (6). Moreover, 30-second epochs may conceal transient sleep stage intrusions and microstructural dynamics, such as frequent stage transitions, which could be clinically relevant. Several studies suggest that important sleep dynamics can occur within shorter time windows than the conventional 30-second epochs allow (7–10).

Automated classification of smaller segments has been explored (10–13), e.g. Cesari et al. (14) applied 5-second scoring in patients with Parkinson’s disease and found increased microstructural sleep instability, i.e. fragmented NREM sleep and short REM episodes, that were not captured in 30-second scoring. Perslev et al. (15) found that analyzing smaller segments than 30-second epochs improved automatic detection of sleep apnea. Such studies suggest that a higher temporal sleep stage resolution may enhance the characterization of sleep architecture and increase diagnostic precision.

U-Sleep, a state-of-the-art deep learning model with flexible scoring resolution, has shown high performance (F1-score was 0.79) in automated scoring of conventional 30-second epochs (15). However, despite the flexibility to score sleep at higher temporal resolutions, it has only been trained on and validated against human-scored 30-second epochs. Previously, our group evaluated the originally published U-Sleep model’s performance in 5-second mini-epochs by comparing its mini-epoch scorings to human-scored 5-second mini-epochs (16). We found significantly more stage transitions when the PSG was assessed in mini-epochs than 30-second epochs, supporting that epochs may conceal important aspects of sleep dynamics. Although the original U-Sleep model was indeed able to classify sleep in shorter segments, its agreement with human-scored mini-epochs was lower than with human-scored epochs (F1-score of 0.54 for mini-epochs vs. 0.78 for epochs) (16). This was likely due to the model’s original epoch-based training (i.e. it was trained to browse through longer intervals and ignore smaller intermittent stage elements if another stage was overall predominant), suggesting that it must be trained on human-scored mini-epochs to obtain a high performance in the mini-epoch setting. However, training an automated classifier to score 5-second mini-epochs at a level comparable to how the original U-Sleep model performs in 30-second epochs would require a substantial number of human-scored mini-epochs, being an unfeasible human workload. For instance, the original U-Sleep model was trained on 19,924 full-night PSGs, comprising millions of 30-second epochs (15).

A potential solution to this challenge is transfer learning, which enables models originally trained on large datasets to be fine-tuned and optimized for specific tasks using limited new data. Transfer learning involves re-training a pre-trained model to adapt it to a new but related task. Ganglberger et al. (17) showed that transfer learning could effectively fine-tune existing sleep scoring models across different datasets, reducing the dependency on large human-scored datasets. Similarly, Guillot et al. (18) applied transfer learning to adapt an automated sleep classifier to diverse PSG montages and demographics suggesting it as a relevant tool for optimizing existing models.

In this present study, by applying transfer learning to U-Sleep, we therefore aim to optimize its performance in 5-second mini-epoch sleep staging using 48,000 human-scored mini-epochs for re-training, validation and testing. Moreover, we compare our human and automatically scored mini-epochs to the conventional human-scored 30-second epochs. Our goal is to facilitate a fast, more detailed, and precise sleep stage analysis by optimizing automated scoring of 5-second mini-epochs.

## Methods

### Cohort

In the period from February 2015 to July 2024, 155 narcolepsy type one (NT1) patients and their 123 non-narcoleptic siblings were recruited to The Norwegian Centre of Expertise for Neurodevelopmental Disorders and Hypersomnias (NevSom), Department of Rare Diagnosis, Oslo University Hospital (OUS). Diagnosis was confirmed in patients and excluded in siblings according to the ICSD-3 criteria (19) by European Sleep research society (ESRS) certified somnologists and NT1 experts (S.K.H and B.H.H). Aiming to include mini-epochs across different health profiles for the optimization of U-Sleep, we randomly selected 100 PSGs from this cohort (38 NT1 patients (age 22.3 ± 8.7 years, 23 females (60.5%)) and 62 non-narcoleptic siblings (age 23.2 ± 12.3 years, 36 females (58.1%))), with no significant difference in age and sex, to be included in the present study. As the selection of individuals was random, the sample included both NT1 patients with and without a non-narcoleptic sibling, and non-narcoleptic siblings with and without a related NT1 patient in the sample. PSG exclusion criteria were a total sleep time (TST) <6 hours and/or apnea-hypopnea index (AHI) >5.

The patients paused their use of stimulants, antidepressants, and/or sodium oxybat for 14 days prior to the PSG. One included patient only paused modafinil nine days before the PSG. Siblings were not required to pause medication. Two patients and five siblings had either allergies, epilepsy, ulcerative colitis, or hypothyroidism and used medications with potential side effects of drowsiness or somnolence reported in 1-10% of users (loratadine, desloratadine, lamotrigine, and mesalamine, levothyroxine), however, the siblings reported no subjective increase in sleepiness and their PSGs were normal.

The study is ethically approved by the Norwegian South-East Regional Committees for Medical and Health Research Ethics (REK) (2014/450), and written informed consent was signed by all individuals or their parents in case of minors <16 years. A subset of the individuals were included in previous studies conducted by our research group (16,20–30).

### Data

The 100 PSGs used for 5-second mini-epoch scorings were recorded using the SOMNOmedics Plus system (DOMINO software, version 2.9.0, SOMNOmedics, Randersacker, Germany), following the recording standards specified by the AASM (1). The PSG data included the following channels: EOG1:A2, EOG2:A1, C3:A2, C4:A1, F3:A2, F4:A1, O1:A2, O2:A1 (sampled at 256 Hz), and two EMG m. submentalis channels (sampled at 256 Hz or 512 Hz). To enhance the chance of covering all sleep stages, two 20-minute segments (240 mini-epochs per segment, i.e. a total of 480 mini-epochs per PSG) were randomly selected from each PSG between the’lights off’ and’lights on’ marks. The 20-minutes segment length was chosen to balance the workload of manual scoring with the need to retain sufficient temporal context for the model, and to align with the segment length of 17.5 minutes used in the original U-Sleep study by Perslev et al. (15). In total, the dataset comprised 48,000 human-scored 5-second mini-epochs, which were all manually scored by an experienced human scorer (J.V., a ESRS certified somnologist-sleep technician).

### Manual mini-epoch scoring procedure

The scoring procedure followed a structured protocol developed specifically for 5-second mini-epochs. An overview of the procedure is shown in Figure 1. Initially, the human scorer spent 10 minutes scrolling in SOMNOmedics DOMINO (the scoring software used in our clinical practice) to get familiar with the PSG data. The human scorer was blinded to age and sex of the individuals, patient/sibling status and furthermore, to the nocturnal timing and the full hypnogram.

**Figure 1:**
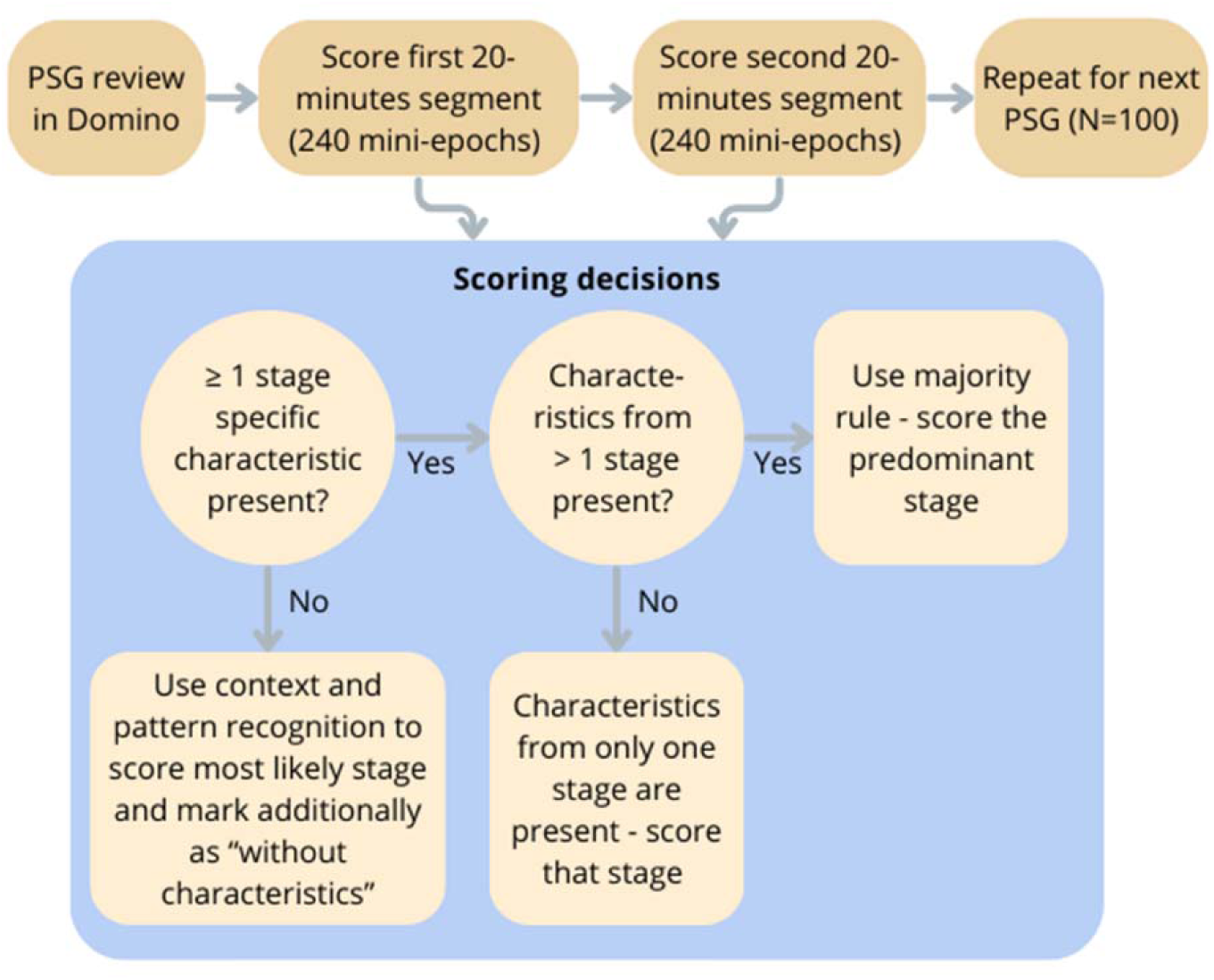
Overview of the manual scoring procedure for 5-second mini-epochs. The human scorer first reviewed the full-night PSG in DOMINO for 10 minutes, then scored two 20-minute segments (240 mini-epochs each) per PSG in MATLAB. The scoring decision for each mini-epoch follows a structured decision tree: If no clear characteristics were present, the most likely stage was determined based on surrounding context and background EEG activity, and the mini-epoch was marked as “without characteristics”. If only stage specific characteristic from a single sleep stage was present, the corresponding stage was scored. If characteristics from multiple stages were present, the predominant stage was scored. This procedure was repeated for all 100 PSGs in the dataset.

The mini-epochs were then shown and scored in a MATLAB 2020b based scoring interface with a setup mimicking the SOMNOmedics DOMINO software layout. The human scorer was allowed to scroll back and forth between the mini-epochs within a 20-minute segment. Each mini-epoch was scored as independently as possible; however, it was allowed to rely on context from previous or subsequent mini-epochs to support decision-making.

### Scoring rules

Our proposed rules for mini-epoch scoring are an adaptation of the AASM rules developed collaboratively in a group of sleep clinicians, including two ESRS certified somnologists (S.K.H. and J.V.), and sleep researchers/technicians (L.F.F., J.A.E.C., R.V., and H.T.J.). This process involved thorough discussions and repeated examinations and reviews of mini-epochs to refine the scoring rules. The full mini-epochs scoring manual is available in Supplementary Material.

The mini-epochs were scored based on the presence of stage-specific characteristics, as summarized in Table 1, exemplified in Figure 1. Notably, if clear characteristics were absent in a mini-epoch, the surrounding context was used to determine the most likely stage and the given mini-epoch (N1, N2, or REM) was additionally marked as “without characteristics”; this was not applicable for wake and N3 (where characteristics are mandatory).

**Table 1:** Stage-specific characteristics used for manual sleep stage scoring in 5-second mini-epochs.

| Stage characteristics | Scoring rules |
| --- | --- |
| **Wake**<br>≥50% Alpha activity | Score wake if alpha activity is present in more than 50% of the mini-epoch. |
| **N1**<br>Slow eye movements<br>Vertex sharp waves | Score N1 if a slow eye movement or vertex sharp wave is present.<br>If characteristics are absent, but the surrounding mini-epochs are scored as N1, and/or the background EEG resembles N1 or does not suggest another stage; score “N1 without characteristics”. |
| **N2**<br>Sleep spindles<br>K-complexes | Score N2 if sleep spindle or K-complex are present. If characteristics are absent, but the surrounding mini-epochs are scored as N2, and/or the background EEG resembles N2 or does not suggest another stage; score “N2 without characteristics”. |
| **N3**<br>High-amplitude, low-frequency delta waves (SWS) | Score N3 only if high-amplitude, low-frequency delta waves (SWS) are present in more than 50% of the mini-epoch. |
| **REM**<br>Rapid eye movements | Score REM if a rapid eye movement is present. If no rapid eye movements are observed, but the surrounding mini-epochs are scored as REM, and/or the background EEG resembles REM or does not suggest another stage; score “REM without characteristics”. |

### Automated mini-epoch scoring procedure

As in our previous smaller study of 5-second mini-epochs (16), we used U-Sleep 2.0, a deep learning model developed by Perslev et al. (15) as the basis for automated sleep stage classification. The U-Sleep model is a fully convolutional neural network that returns a hypnogram based on a single EEG/EOG combination. If data from more channels are present in the PSG, the hypnogram is determined by majority voting from all possible EEG/EOG combinations. U-Sleep pre-process the input data by resampling the signals to 128 Hz and removing outliers of the signals.

In the present study, U-Sleep predicted sleep stages in each 20-minutes segment, using the six EEG channels; C3:A2, C4:A1, F3:A2, F4:A1, O1:A2, and O2:A1 and the EOG channels EOG1:A2 and EOG2:A1. Application of the original U-Sleep model per segment created a baseline mini-epoch scoring.

The source code for the automated U-Sleep model (15) was obtained via a link available in the original article (https://github.com/perslev/U-Time, downloaded March 29, 2023) and the weights for the pre-trained neural network were kindly made available by the first author on request.

### Transfer learning

After our U-Sleep baseline mini-epoch scoring, we then optimized the original U-Sleep model via transfer learning on 20-minutes segments of human-scored 5 second mini-epochs, since the original U-Sleep model was trained on 17.5 minutes segments so as to at least provide the model with a similar duration of input (15). The human scored PSG segments were then divided into three sets to be used for training (n=80), validation (n=10), and testing (n=10) (Table 2), ensuring that both 20-minutes segments from each PSG remained within the same set. This prevented data leakage between training and testing. The distribution of NT1 status, age, and sex was maintained across the sets. We modified the model to train on and predict sleep stages every 5 seconds, and fine-tuned all components (encoder, decoder, and segment classifier) with a learning rate of 10^-7^ using an Adam optimizer and a cross-entropy loss function. Convergence was monitored using the F1-score on the validation set, and model training was terminated after 500 consecutive mini-epochs with no apparent increase in validation F1. The code was implemented in Tensorflow version 2.8 (31,32) based on the original source code.

**Table 2:** Number of PSGs and mini-epochs in the training, validations and test set used to optimize and evaluate the U-Sleep model.

| **Set** | **PSGs (n)** | **Mini-epochs (n)** |
| --- | --- | --- |
| Training | 80 | 38400 |
| Validation | 10 | 4800 |
| Test | 10 | 4800 |
| **Grand total** | 100 | 48000 |

### Statistical analysis

To assess inter-scorer agreement, confusion matrices were constructed for each pairwise scorer combination (human-scored epochs vs. human-scored mini-epochs, U-Sleep vs. human-scored epochs, and U-Sleep vs. human-scored mini-epochs). Mini-epochs labeled with artifacts (i.e., that could not be scored) by the human scorer (221/48,000 mini-epochs, 0.46%) were removed from the analysis before constructing the confusion matrices. As a result, the total number of observations in each segment’s confusion matrix reflects only the artifact-free mini-epochs.

To reduce random variation and mitigate the problem that the 20-minute segments occasionally lacked one or more sleep stages, resulting in missing agreement estimates, the two 20-minute segments from each PSG were combined to generate a single confusion matrix per PSG. This ensured that stage-wise agreement estimates were based on a more complete dataset across PSGs. The F1-score, i.e., harmonic mean of precision and recall, ranging from 0 (no agreement) to 1 (perfect agreement), was then calculated for each sleep stage and overall, i.e. across all stages, per PSG directly from the confusion matrices using the standard formula:

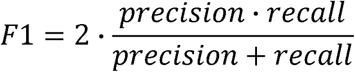

where *precision* = *TP*/(*TP* + *FP*) and recall = *TP*/(*TP* + *FN*) were computed for each sleep stage. *TP*, *FP* and *FN* denote true positives, false positives and false negatives, respectively. To account for class imbalance, each stage’s F1-score was weighted according to its prevalence in the dataset when calculating the mean F1-score across all stages.

Since one or more sleep stages sometimes were absent in the combined two 20-minutes of PSG data, missing values should be handled carefully. For stages that were absent in a given PSG in both the truth and prediction (TP=0 and FP=0), the corresponding F1-score was zero, representing a missing agreement value rather than true disagreement. To avoid artificially lowering mean values, these zeros were excluded from stage-wise averages. The number of PSGs contributing to each mean value is reported in the results.

Bootstrapping (n=1,000) was used to estimate means and 95% confidence intervals for F1-scores and sleep stage fractions as these measures were not normally distributed. For statistical comparisons between scoring methods, the non-parametric Wilcoxon signed rank-sum test was applied to compare F1-scores, sleep stage fractions and stage transitions across PSGs.

The following scorings were compared pairwise: Human-scored 5-second mini-epochs; conventional human-scored 30-second epochs (splitting each 30-second epoch into six 5-second mini-epochs and assigning the epoch’s label to each mini-epoch); automated 5-second mini-epochs scorings by the original U-Sleep model (15); automated 5-second mini-epochs scorings by the optimized U-Sleep model.

Confusion matrices were for visualization purposes either row-or column-normalized depending on the evaluation objective. When comparing two human scorings (e.g., epochs vs. mini-epochs), the confusion matrix was precision-weighted (column-normalized) to reflect scorer alignment. When evaluating automated vs. human scorings, matrices were recall-weighted (row-normalized) to reflect performance across sleep stages.

## Results

### Mini-epochs versus epochs

To contextualize the 48,000 human-scored mini-epochs, we first compared them to the conventional human-scored 30-second epochs. We examined differences in sleep stage distribution and classification to assess how increased temporal resolution affects sleep staging.

Figure 2a shows distribution of sleep stages across PSGs in mini-epochs and epochs. Though the scored segments only represent a selection of the full-night PSG recordings, the fractions were similar to typical full-night sleep stage fractions (33) and followed the typical sleep distribution, with N2 being the dominant stage and wake and N1 appearing less often. However, mini-epoch scorings contained significantly more wake, N1 and N2, and less N3 and REM sleep than the epochs (all p<0.001). Furthermore, we analyzed sleep stage transitions per PSG. Analyzing mini-epochs resulted in 1.70 ± 0.79 stage transitions per minute while epoch analysis resulted in significantly less stage transitions, i.e. only 0.21 ± 0.12 per minute (p<0.001), reflecting that the higher temporal resolution enables detection of frequent stage shifts not captured by epochs.

**Figure 2:**
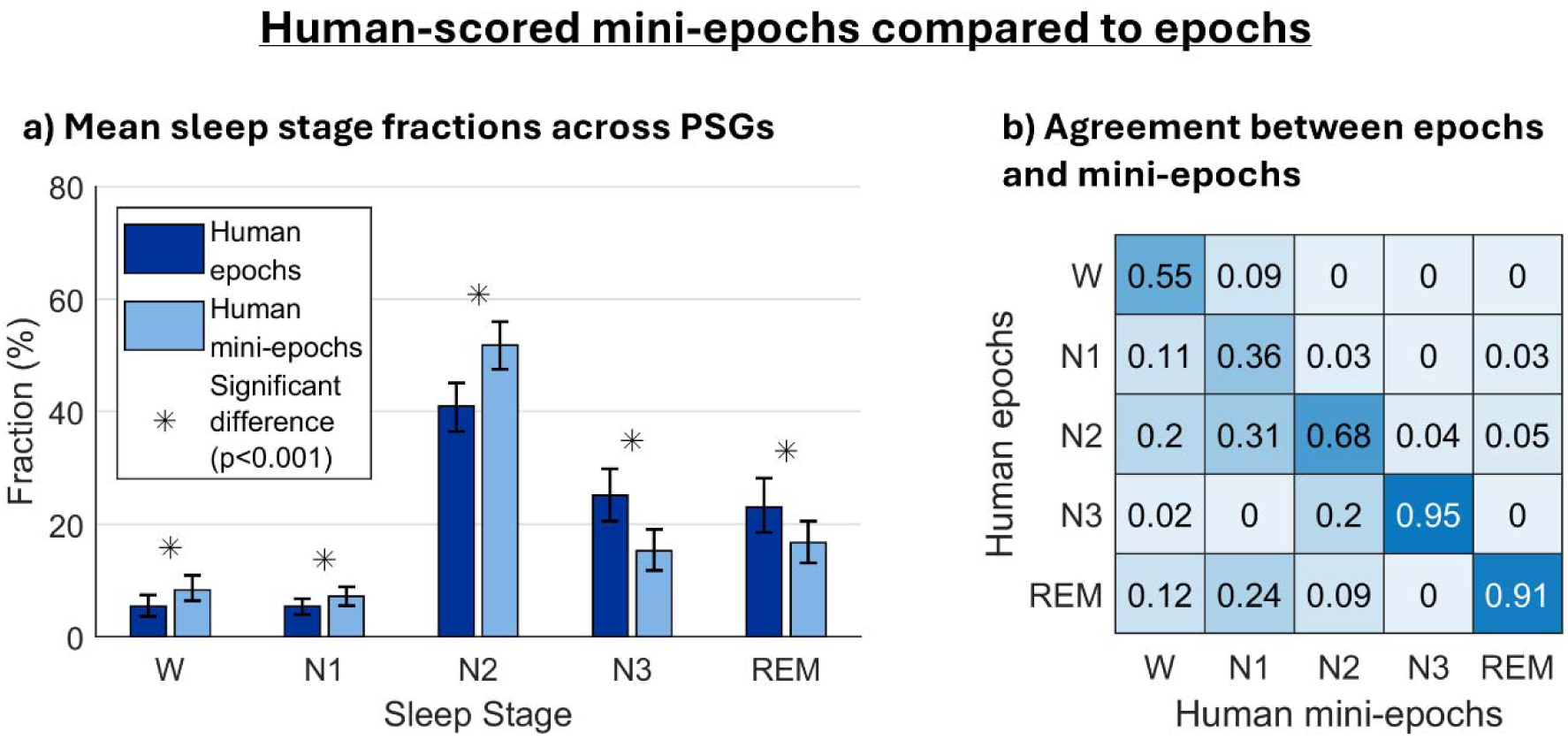
The figure is based on 100 PSGs, each contributing two 20-minute PSG segments. (a) Mean fraction of each sleep stage (W, N1, N2, N3, REM) in human scored 30-second epochs and human-scored mini-epochs. The bar heights show the means across PSGs, and the whiskers represent 95% confidence intervals obtained via bootstrapping. Asterisks indicate significant differences (p<0.001) obtained with the signed Wilcoxon test. (b) Precision weighted, column-wise normalized confusion matrix between human-scored epochs (rows) and human-scored mini-epochs (columns). Diagonal values represent agreement for each sleep stage. Off-diagonal values show misclassification patterns (e.g., row 3, column 2 indicates that 31% of the mini-epochs scored as N1 in the human mini-epoch scorings comes from N2 in the human-scored epochs).

Figure 2b visualizes the confusion matrix between mini-epochs and epochs. The rows represent the sleep stages scored in the epochs while the columns represent the corresponding scorings in the mini-epochs. The matrix was column-wise normalized (precision-weighted), meaning that each value represents the proportion of times a given mini-epoch classification originated from a specific epoch sleep stage. As seen, the agreement between epochs and mini-epochs, presented in the diagonal of the matrix, was highest in N3 and REM sleep, and lowest in N1 sleep.

Of the stage wake-scored mini-epochs, 20% came from N2 epochs whereas 11% and 12% came from N1 and REM sleep epochs, respectively. Of the N1-scored mini-epochs, 31% came from N2 epochs and 24% came from REM sleep epochs. Regarding the N2 scored mini-epochs, 20% came from N3 scored epochs, whereas 9% came from REM sleep epochs. N3 mini-epochs mainly came from N3 epochs, with only 4% coming from N2 epochs. Of the REM scored mini-epochs, 3% came from N1, and 5% from N2 scored epochs.

### U-Sleep’s performance in the test-set before and after transfer learning

To assess the effect of optimizing U-Sleep via transfer learning, we compared predictions from both the original U-Sleep model, and the transfer learning optimized U-Sleep model to the human-scored mini-epochs in the test set (20 segments from 10 PSGs not used in model training or validation, 4,800 mini-epochs in total).

Figure 3 summarizes the mean performance of U-Sleep in mini-epochs before and after transfer learning. Figure 3a shows stage-wise and overall performance of U-Sleep in terms of the F1-scores. Transfer learning improved U-Sleep’s performance across all stages except REM, where performance decreased slightly. The overall F1-scores across PSGs improved significantly from 0.74 to 0.81 (p<0.05). A significant improvement was also observed in stage wake (p<0.05), while changes in other stages did not reach statistical significance. Although the increase in N1 was visibly large, the difference was not statistically significant, most likely due to high F1-score variability across PSGs in N1.

**Figure 3:**
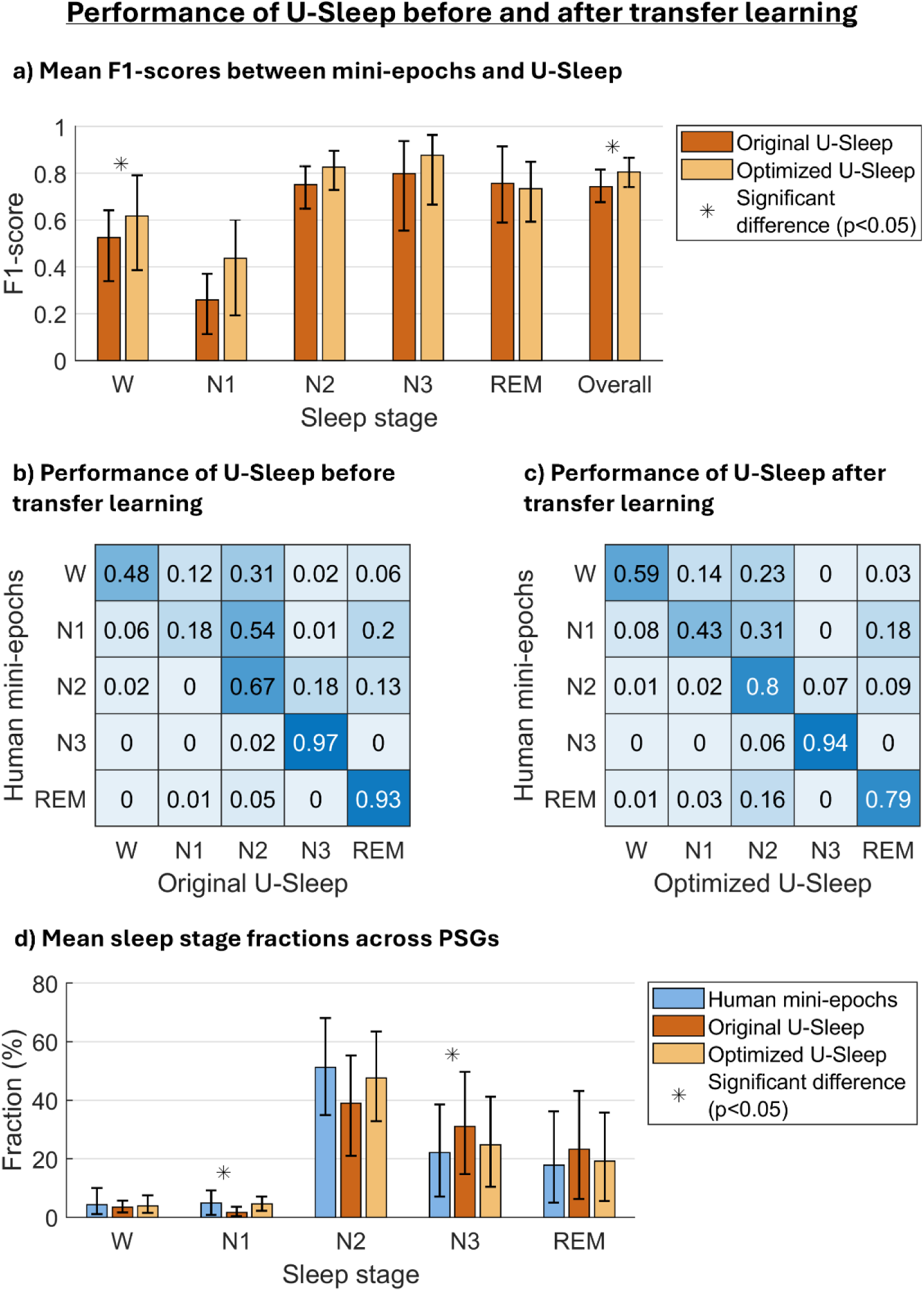
The figure is based on the test data from 10 PSGs each contributing two times 20-minutes of PSG data. (a) Performance of U-Sleep before and after transfer learning in terms of the F1-score per stage and overall (i.e. the weighted mean F1-score across all stages) when compared to the human-scored mini-epochs. Mean F1-scores per stage were calculated across PSGs with at least one mini-epoch in the given stage (scored by either the human scorer or the model). Included PSGs per mean: human vs original U-Sleep model - wake (n=10), N1 (n=8), N2 (n=9), N3 (n=8), REM (n=6); human vs optimized U-Sleep model - wake (n=10), N1 (n=10), N2 (n=10), N3 (n=8), REM (n=6). (b-c) Recall-weighted, row-wise normalized confusion matrices between human-scored mini-epochs vs (b) original and (c) optimized U-Sleep model. Each cell shows the fraction of mini-epochs scored in a given stage by U-Sleep (columns), conditional on the human mini-epoch scorings (rows). Diagonal values represent agreement per stage and off-diagonal values show misclassifications (e.g. row 2, column 3 shows that U-Sleep scored N2 in 54% (original model) and 31% (optimized model) of the mini-epochs scored as N1 in human-score miniepochs). (d) Mean sleep stage fractions scored by human, original U-Sleep, and optimized U-Sleep. In (a) and (d) the bar heights show the means across PSGs, and the whiskers show bootstrap-derived 95% confidence intervals. Asterisks denote significant differences (p<0.05) obtained with the signed Wilcoxon test.

To further evaluate the scoring discrepancies, recall-weighted confusion matrices were constructed with human-scored mini-epochs as the ground truth versus U-Sleep before transfer learning (original U-Sleep), and U-Sleep after transfer learning (optimized U-Sleep), respectively, as predictions (Figures 3b-c). Diagonal values indicate stage-wise agreement, which improved after transfer learning for wake, N1, and N2, at the cost of REM agreement. Notably, stage REM, N1, and N2 included mini-epochs both with and without stage-specific characteristics. As specified in Figure 5, the reduced REM agreement was primarily driven by mini-epochs without stagespecific characteristics i.e. mini-epochs resembling stage REM EEG-wise (and possibly EMG-wise) but lacking rapid eye movements.

After transfer learning, U-Sleep’s most frequent misclassifications occurred between N1 and N2. Of the human-scored N1 mini-epochs, 31% were predicted as N2 and 18% as REM by the optimized U-Sleep model. For human-scored wake mini-epochs, 23% were predicted as N2 and 14% as N1. Human-scored REM mini-epochs were predicted as N2 in 16% of the cases.

Figure 3d shows the sleep stage distributions based on mini-epoch scorings by the human scorer, the original U-Sleep model and the optimized U-Sleep model, respectively. Before transfer learning, the original U-Sleep model significantly underestimated N1 and overestimated N3 compared to the human-scored mini-epochs, however, these differences were no longer present after model optimization.

In terms of sleep stage shifting, the number of stage transitions differed significantly between both U-Sleep models and the human-scored mini-epochs. Analyzing human-scored mini-epochs resulted in an average of 1.70 ± 0.86 transitions per minute, whereas the original U-Sleep model produced fewer transitions (1.06 ± 0.99, p < 0.05). The optimized U-Sleep model also produced fewer transitions compared to the human-scored mini-epochs (1.34 ± 0.80, p < 0.05), however, the number of transitions was significantly higher than the original U-Sleep model (p < 0.05), reflecting an improved ability to capture sleep fragmentation after transfer learning.

In Figure 4, the mean overall F1-scores are shown for all pairwise comparisons between human-scored epochs, human-scored mini-epochs, and U-Sleep scorings before and after transfer learning, respectively, in the test set. Before transfer learning, the original U-Sleep model agreed more with the conventional epoch scoring than with the mini-epoch scorings (F1=0.85 vs. 0.74), reflecting that the model was originally trained on epochs. After transfer learning, the optimized U-Sleep model’s agreement with epochs decreased (F1=0.81), while agreement with mini-epochs increased significantly (F1=0.81, p<0.05).

**Figure 4:**
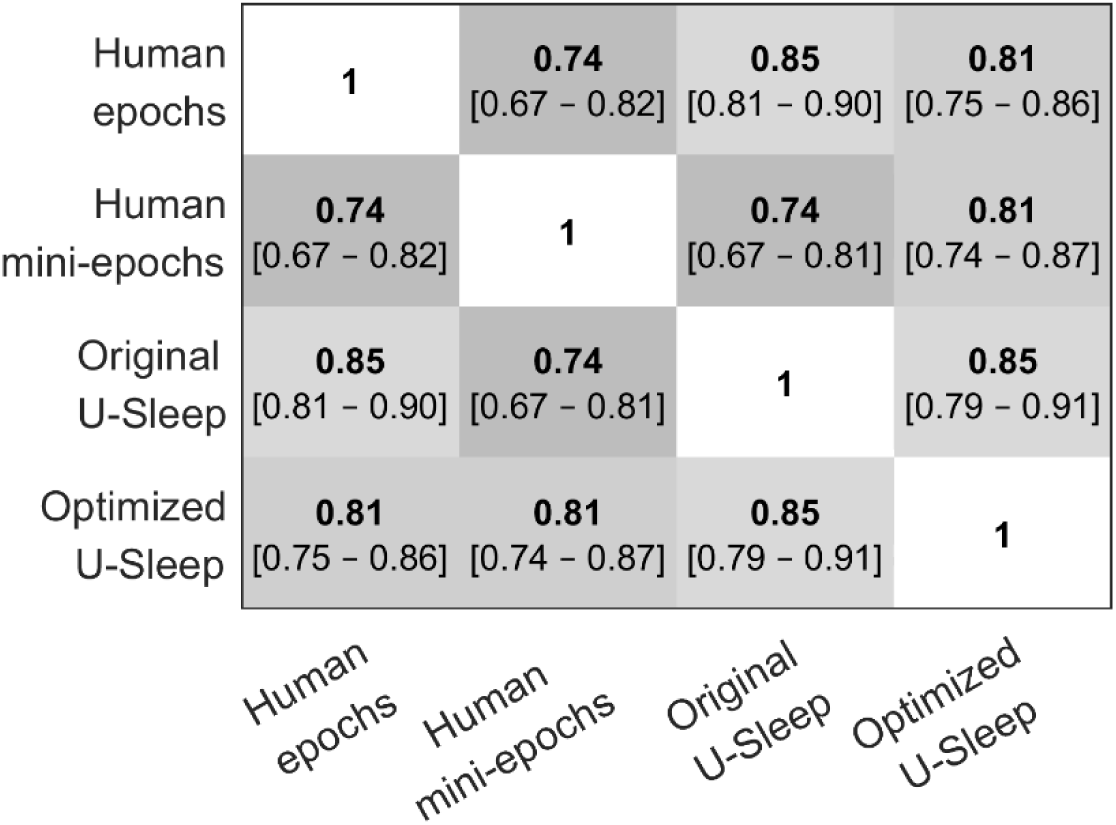
Pairwise agreement rates between conventional epochs, human-scored mini-epochs, and U-Sleep before and after transfer learning across the 10 PSGs in the test set represented as mean overall F1-scores (i.e. across all sleep stages) and bootstrapped 95% confidence interval.

**Figure 5:**
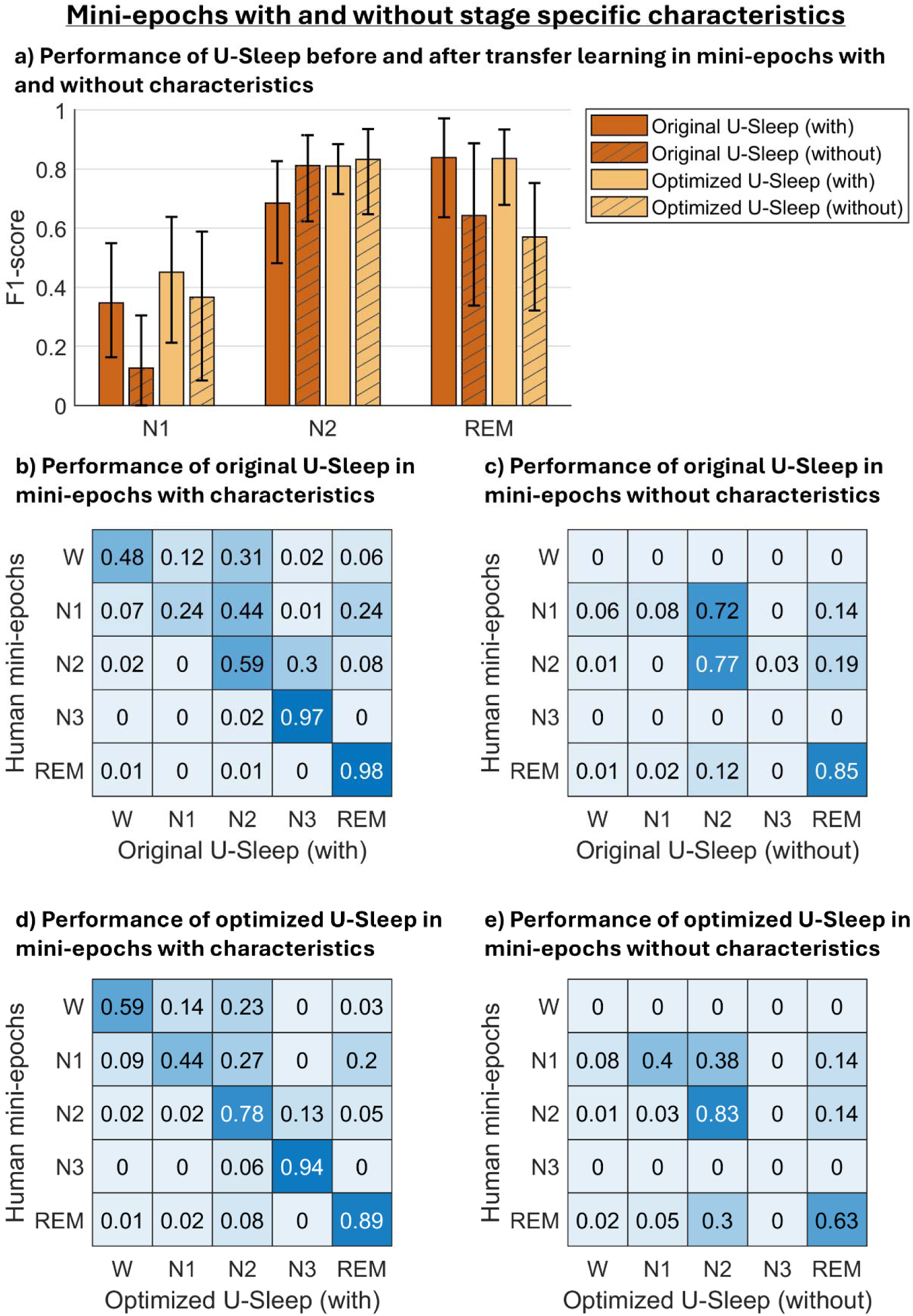
The figure is based on test set data from 10 PSGs, each contributing two 20-minutes PSG segments. (a) F1-scores between human-scored mini-epochs and U-Sleep before and after transfer learning, stratified by whether the mini-epochs included stage-specific characteristics or not according to the human scorer. Note that the option to mark a mini-epoch as “without characteristics” did not apply for wake and N3. Due to limited data points, statistical comparisons were underpowered and formal significance testing was not performed. Bar heights show the mean F1-score across PSGs for N1, N2, and REM, with whiskers representing 95% confidence intervals obtained via bootstrapping. (b–e) Recall-weighted, row-wise normalized confusion matrices between human-scored mini-epochs and U-Sleep, computed separately for mini-epochs with (b, d) and without (c, e) stage-specific characteristics. Panels (b–c) show the performance of the original U-Sleep model; (d–e) show the performance of the optimized model. Each matrix cell represents the fraction of mini-epochs classified as a given stage by U-Sleep (columns), conditional of the human mini-epoch scoring (rows). Diagonal values reflect agreement per stage, while offdiagonal values represent misclassifications.

### U-Sleep’s performance in mini-epochs with and without stage-specific characteristics

To investigate how U-Sleep performed in mini-epochs without clear stage-specific characteristics, we stratified the mini-epoch dataset based on whether the human scorer had marked the mini-epochs as “without characteristics” or not. On average, 30.8±16.6% of the mini-epochs were marked as “without characteristics”. Specifically, 38% of N1 and 43.9% of N2 were marked without characteristics, and 37.7% of REM mini-epochs lacked eye movements.

Figure 5 presents an analysis of performance of the original and optimized U-Sleep models across mini-epochs with and without characteristics in the test set. As shown in Figure 5a, in N1 and REM, U-Sleep agreed more with human-scored mini-epochs with characteristics than mini-epochs without characteristics, both before and after transfer learning (e.g. after transfer learning the F1-scores were 0.45 versus 0.37 for N1 with versus without characteristics, respectively, and 0.84 versus 0.56 for REM with versus without characteristics, respectively). For N2, the opposite pattern was observed for U-Sleep both before and after transfer learning, although the difference was minimal after transfer learning (F1-scores were 0.81 with versus 0.83 without characteristics after transfer learning). We did not perform significance testing in this analysis because the limited number of data points made statistical comparisons underpowered.

The recall-weighted confusion matrices in Figures 5b–e support these findings. The most common misclassifications in mini-epochs without characteristics occurred between N1 and N2, but these decreased notably after transfer learning.

## Discussion

In this study, we investigated whether U-Sleep, existing automated sleep stage classifier (15) with high performance in 30-second epochs, could be optimized via transfer learning to more accurately score sleep in 5-second mini-epochs. Using a dataset of 48,000 human-scored mini-epochs following AASM-inspired scoring rules, we demonstrated that transfer learning significantly improved the model’s agreement with human scoring and achieved a high performance with an overall F1-score of 0.81, supporting the feasibility of automated mini-epoch sleep staging.

Before assessing model performance, we compared our human-scored mini-epochs to conventional epoch scorings to understand how increased temporal resolution affects sleep stage distribution and structure. The same human (J.V.) scored both the mini-epochs and the epochs, which may have introduced a scoring bias although done with minimum one year in between. However, this approach was intentionally chosen to avoid disagreement between mini-epochs and epochs reflecting differences between scorers. We found that the numbers of stage transitions were significantly higher in the human-scored mini-epochs than epochs, showing that mini-epochs provide a more detailed representation of sleep architecture as also suggested by our previous smaller study (16). Moreover, mini-epoch scoring resulted in significantly more wake, N1, and N2, and less N3 and REM compared to conventional 30-second epoch scoring. This difference might partly be explained by the AASM epoch scoring rule, where ≥20% of an epoch with N3 activity results in the entire epoch being scored as N3. When N3 appears together with another stage, the non-dominant stage (typically N2) is ignored in epoch scoring because the ≥20% rule favors N3. However, this non-dominant stage can be captured in mini-epoch scoring likely contributing to the higher N2 and lower N3 proportions observed in mini-epochs. The reduced REM sleep proportion in mini-epochs may be due to our exclusion of low muscle tone as a REM-specific characteristic, causing some ambiguous REM-like mini-epochs (i.e. REM-like mini-epochs without rapid eye movements) to be scored as N1 or N2. This was, however, chosen because low muscle tone was also observed in mini-epochs with clear NREM or W characteristics, i.e. we concluded that low muscle tone could not be used as a REM-specific feature in mini-epochs without rapid eye movements. It remains unknown how the sleep stage distributions would look if we had included low muscle tone as a REM-specific characteristic.

A key advantage of mini-epochs is the ability to capture frequent stage transitions and brief events that are missed in epochs. For example, brief intrusions of wake or sleep may appear only for a few seconds and would be averaged out in traditional epoch scoring. Previous studies have shown that repeated MSLT tests in the same patient can give different results and even lead to change in diagnoses (34,35). Although this could reflect changes in disease state (progression, remission), the known problems with the 30-second epoch length (several sleep stages in one epoch; interrater variability, etc.) are also likely to contribute. Hence, such diagnostic instability underlines the need for more precise and physiologically sensitive sleep analysis methods. Mini-epochs could potentially contribute with new disease biomarker, and play a role in improving diagnostic accuracy, particularly in disorders like narcolepsy and it borderland, where frequent stage transitions are common and may be missed in traditional scoring frameworks (36).

However, higher temporal resolutions also come with trade-offs. By requiring a sleep stage label for each 5-second period, mini-epochs increase the number of decisions, which can be difficult particularly in biologically gradual transition periods, possibly increasing inter-rater variability. On the other hand, shorter intervals might reduce the well-known problem of epochs containing characteristics from multiple stages. Younes et al. (6) reported that such “equivocal epochs” are the main contributors to epoch scoring disagreement and that equivocal epochs often result in nearly random scoring. Mini-epochs help reduce this problem by using shorter time intervals, which means that each mini-epoch is less likely to include characteristics from more than one sleep stage. This was also reported by the human scorer in present study. Whether inter-rater variability in human mini-epoch scoring will increase or decrease compared to conventional epochs remains to be evaluated in future studies.

Despite the different advantages, human scoring of mini-epochs is far too time-consuming to be feasible in clinical settings. To address this limitation, we applied an automated solution by using our human-scored mini-epoch dataset as ground truth for optimizing the U-Sleep model via transfer learning. We specifically chose the U-Sleep model due to its previously shown high performance (15,37), however in 30-second epochs, and its flexibility to handle different temporal resolutions.

In our test set, optimizing U-Sleep via transfer learning led to a significant increase in the overall F1-score from 0.74 to 0.81 when compared to human-scored mini-epochs. Hence, the optimized U-Sleep modeĺs overall score falls within the range reported in previous studies of automated sleep staging (F1=0.70–0.87) (15,38–41), though in epochs, and is notably higher than the F1-score observed in our previous study of the original U-Sleep model’s performance in human-scored mini-epochs (F1=0.54) (16). Our present stage-wise results also showed trends of improvement, particularly in wake and N1, although only wake reached statistical significance. The lowest performance of U-Sleep was observed in N1, consistent with previous studies (in epochs) reporting that automated sleep stage classifiers generally perform worst in N1 (11,15,17,38,42,43). N1 is also the stage with greatest human inter-rater variability (6,44,45). The highest optimized U-Sleep model performance was achieved in N3, which has likewise previously been reported as the stage in which automated classifiers performs best (46), although these findings were based on 30-second epochs.

The relatively lower agreement in REM sleep after U-Sleep optimization via transfer learning likely reflects both methodological and biological challenges. Our mini-epoch scoring rules (Supplementary Material) deviated from the AASM standard as we (as explained above) felt the need to exclude low muscle tone as a REM-specific characteristic potentially making stage REM more difficult to score consistently. This possibly required U-Sleep to learn a new representation of REM which may have increased the learning burden, particularly if our human scorings of ambiguous REM mini-epochs were not fully consistent.

Additionally, many REM-scored mini-epochs lacked rapid eye movements and often resembled light NREM sleep, especially N2 (if mini-epochs also lacked spindles and K-complexes). The optimized U-Sleep model frequently misclassified such mini-epochs as N2 (30%) which may reflect true biological ambiguity or mixed-stage phenomena rather than misclassification.

While transfer learning improved U-Sleep’s overall performance in mini-epochs (F1=0.81), this performance did not exceed the performance of the optimized U-Sleep model in epochs (F1=0.81). This may be due to several factors. First, Perslev et al. (15) trained the original U-Sleep model on large-scale 30-second epoch data which has sustained influence on its predictions in our present study even after transfer learning. For instance, the original U-Sleep model’s scorings resulted in the fewest stage transitions per segment, while the optimized U-Sleep model’s scorings resulted in more transitions, but still fewer than the human-scored mini-epochs. This suggests that though the optimized U-Sleep model did become much better at scoring mini-epochs and reached a high performance, its predictions remained smoother (i.e., with fewer transitions between stages).

Second, both U-Sleep and the human mini-epoch scorings may contain frequent transitions, but if these are not exactly aligned, the scorings will disagree. Even small temporal mismatches such as one transition occurring one mini-epoch earlier or later will count as disagreement. In contrast, epoch-based scoring inherently averages over brief sleep or wake intrusions, which may lead to better agreement—even if the underlying biologically true stage transitions are not perfectly captured.

Third, some of the disagreement between human and automated scorings of mini-epochs might reflect true uncertainty in the data. For example, the boundaries between N1 and N2 sleep are biologically gradual and sometimes ambiguous, even to expert human scorers (44). This ambiguity can limit the achievable agreement, not because the model performs poorly, but because the underlying signal lacks a clear categorical boundary. So, even a perfect algorithm might struggle to agree with a human scorer on these transitional periods, just as two trained human scorers can disagree (44,45). In fact, studies reporting inter-rater agreement for human scorers using the F1-score have found values ranging between 0.70 and 0.79 (15,38,41)

In average, 30.8% of the human-scored mini-epochs in the present study were marked as lacking defining stage-specific characteristics, making them difficult to score - a problem likely less common in 30-second epochs, where the longer time windows provide more contextual information. These ambiguous mini-epochs without characteristics contribute to uncertainty and may potentially lead to variability in both human and automated sleep stage scoring. We therefore explored how the presence or absence of stage-specific characteristics in mini-epochs influenced U-Sleep’s performance. As expected, agreement between U-Sleep and the human scorer was higher in mini-epochs with characteristics for N1 and REM, both before and after U-Sleep optimization, underlining the model’s reliance on distinct visual characteristics such as slow or rapid eye movements. Interestingly, for N2, U-Sleep performed better in mini-epochs without characteristics, both before and after transfer learning. This counterintuitive result may reflect the model’s ability to detect underlying background patterns associated with N2 even when spindles or K-complexes are absent. However, many of the human-scored N1 mini-epochs without characteristics were also by U-Sleep scored as N2 (72% by the original U-Sleep model and 38% by the optimized U-Sleep model). This indicates a tendency of the U-Sleep model to default to N2 when clear characteristics are absent possibly due to the high prevalence of N2 in both the original epoch-based training dataset and our mini-epoch training data. Together, these findings highlight the trade-off of mini-epoch scoring: it reduces equivocal cases with mixed characteristics but introduces more stage decisions where clear defining characteristics are missing.

We acknowledge that our study has limitations. Transfer learning and evaluation of U-Sleep were performed within one study population, hence external validation is needed to assess generalizability. We did not investigate nor consider human inter-rater variability in mini-epochs in this study; however, this should be explored further in larger studies. Statistical comparisons were limited by relatively small numbers of mini-epochs per stage in the test set which potentially affected the stage-wise F1-scores.

Additionally, we evaluated model performance on 20-minute segments rather than full-night PSGs in which performance remains unknown. While we followed the original U-Sleep approach using 17.5-minute training windows, we did not investigate whether training on shorter windows could influence performance. Mini-epoch were scored without knowing the nocturnal timing, which is a deviation from standard clinical scoring procedures that may have influenced scoring decisions in unknown ways.

Finally, mini-epochs may challenge existing diagnostic frameworks as features such as, for example REM-latency or sleep onset REM (SOREM) criteria are based on epochs and may not translate directly to high-resolution mini-epoch scoring. Traditional diagnostic criteria may need to be reconsidered or adapted if in future applied to mini-epochs.

This study demonstrated that automated sleep staging at high temporal resolution is both feasible and accurate. The mini-epoch framework enabled a more detailed representation of sleep architecture, capturing significantly more stage transitions than in traditional epoch-based scoring. By optimizing the deep learning model U-Sleep on 5-second mini-epochs using human-scored sleep stage data, we achieved a significant improvement in agreement between the optimized model and the human scorer compared to the original model. The performance of the optimized U-Sleep model reached levels equally high in mini-epochs as those previously published of other automated classifiers and as agreement between human scorers in 30-second epochs.

## Acknowledgement

We especially thank all the patients and siblings of our study. This study was funded by a grant from the South-Eastern Norway Regional Health Authority (R.V. grant: 2017070; H.T.J. grant: 2019032), the Norwegian Ministry of Health and Care Services (L.F.F., R.V., J.V., H.T.J., J.A.E.C., S.K.H.), the Lundbeck Foundation (A.N.Z. grant: R347-2020-2439), and The Independent Research Fund Denmark through the project „U-Sleep” (M.P. project number: 9131-00099B).

## References

1. Schulz H. Rethinking sleep analysis. J Clin Sleep Med. 2008;4(2):99–103.

2. Rechtschaffen A, Kales A. A manual of standardized terminology, technics and scoring system for sleep stages of human subjects. Wash DC Gov Print Off. 1968;

3. Iber C, Ancoli-Israel S, Chesson AL, Quan SF. The AASM Manual for the Scoring of Sleep and Associated Events. 2007;

4. Grigg-Damberger MM. The AASM scoring manual: A critical appraisal. Curr Opin Pulm Med. 2009;15(6):540–9.

5. Grigg-Damberger MM. The AASM scoring manual four years later. J Clin Sleep Med. 2012;8(3):323–32.

6. Younes M, Raneri J, Hanly P. Staging sleep in polysomnograms: Analysis of inter-scorer variability. J Clin Sleep Med. 2016;12(6):885–94.

7. Moul DE, Germain A, Cashmere JD, Quigley M, Miewald JM, Buysse DJ. Examining initial sleep onset in primary insomnia: A case-control study using 4-second epochs. J Clin Sleep Med. 2007;3(5):479–88.

8. Younes M, Ostrowski M, Soiferman M, Younes H, Younes M, Raneri J, et al. Odds Ratio Product of Sleep EEG as a Continuous Measure of Sleep State. Sleep. 2015;38(4):641–54.

9. Tanaka H, Hayashi M, Hori T. Statistical features of hypnagogic EEG measured by a new scoring system. Sleep. 1996;19(9):731–8.

10. Koch H, Jennum P, Christensen JAE. Automatic sleep classification using adaptive segmentation reveals an increased number of rapid eye movement sleep transitions. J Sleep Res. 2019;28(2):1–12.

11. Olesen AN, Jørgen Jennum P, Mignot E, Sorensen HBD. Automatic sleep stage classification with deep residual networks in a mixed-cohort setting. Sleep. 2021;44(1):1–12.

12. Aboalayon KAI, Faezipour M, Almuhammadi WS, Moslehpour S. Sleep stage classification using EEG signal analysis: A comprehensive survey and new investigation. Entropy. 2016;18:1–31.

13. Stephansen JB, Olesen AN, Olsen M, Ambati A, Leary EB, Moore HE, et al. Neural network analysis of sleep stages enables efficient diagnosis of narcolepsy. Nat Commun. 2018;9(1):1–15.

14. Cesari M, Christensen JAE, Muntean M lucia, Mollenhauer B, Sorensen HBD, Trenkwalder C, et al. A data-driven system to identify REM sleep behavior disorder and to predict its progression from the prodromal stage in Parkinson’ s disease. Sleep Med. 2021;77:238–48.

15. Perslev M, Darkner S, Kempfner L, Nikolic M, Jennum PJ, Igel C. U-Sleep: resilient highfrequency sleep staging. Npj Digit Med. 2021;4(72):1–12.

16. Follin L, Neergaard A, Viste R, Vevelstad J, Kaufmann T, Ramm-pettersen A, et al. An inter-rater variability study between human and automatic scorers in 5-s mini-epochs of sleep. Sleep Med. 2025;128(October 2024):139–50.

17. Ganglberger W, Nasiri S, Sun H, Kim S, Shin C, Westover MB, et al. Refining sleep staging accuracy: Transfer learning coupled with scorability models. Sleep. 2024;(August):1–11.

18. Guillot A, Thorey V. RobustSleepNet: Transfer Learning for Automated Sleep Staging at Scale. IEEE Trans Neural Syst Rehabil Eng. 2021;29:1441–51.

19. American Academy of Sleep Medicine. International classification of sleep disorders (ICSD), 3rd edition. 2014;

20. Viste R, Soosai J, Vikin T, Thorsby PM, Nilsen KB, Knudsen S. Long-term improvement after combined immunomodulation in early post-H1N1 vaccination narcolepsy. Neurol Neuroimmunol NeuroInflammation. 2017;4(5):1–3.

21. Juvodden HT, Alnæs D, Lund MJ, Agartz I, Andreassen OA, Dietrichs E, et al. Widespread white matter changes in post-H1N1 patients with narcolepsy type 1 and first-degree relatives. Sleep. 2018;41(10):1–11.

22. Juvodden HT, Alnæs D, Lund MJ, DIetrichs E, Thorsby PM, Westlye LT, et al. Hypocretin-deficient narcolepsy patients have abnormal brain activation during humor processing. Sleep. 2019;42(7):1–11.

23. Nordstrand SEH, Hansen BH, Rootwelt T, Karlsen TI, Swanson D, Nilsen KB, et al. Psychiatric symptoms in patients with post-H1N1 narcolepsy type 1 in Norway. Sleep. 2019;42(4):1–9.

24. Nordstrand SEH, Juvodden HT, Viste R, Rootwelt T, Karlsen T ivar I, Thorsby PM, et al. Obesity and other medical comorbidities among NT1 patients after the Norwegian H1N1 influenza epidemic and vaccination campaign. Sleep. 2020;43(5):1–8.

25. Viste R, Lie BA, Viken MK, Rootwelt T. Narcolepsy type 1 patients have lower levels of effector memory CD4+ T cells compared to their siblings when controlling for H1N1-(Pandemrix^TM^)-vaccination and HLA DQB1*06:02 status. Sleep Med. 2021;85:271–9.

26. Viste R, Viken MK, Lie BA, Juvodden HT, Nordstrand SEH, Thorsby PM, et al. High nocturnal sleep fragmentation is associated with low T lymphocyte P2Y 11 protein levels in narcolepsy type 1. Sleep. 2021;44(8):1–12.

27. Juvodden HT, Alnæs D, Lund MJ, Agartz I, Andreassen OiA, Server A, et al. Larger hypothalamic volume in narcolepsy type 1. Sleep. 2023;46(11):1–10.

28. Viste R, Follin LF, Kornum BR, Lie BA, Viken MK, Thorsby PM, et al. Increased muscle activity during sleep and more RBD symptoms in H1N1-(Pandemrix)-vaccinated narcolepsy type 1 patients compared with their non-narcoleptic siblings. Sleep. 2023;46(3):1–14.

29. Hansen BH, Norsted H, Gjesvik J, Thorsby PM, Naerland T, Knudsen-heier S. Associations between psychiatric comorbid disorders and executive dysfunctions in hypocretin-1 de fi cient pediatric narcolepsy type1 Behavior Rating Inventory of Executive Function. Sleep Med. 2023;109:149–57.

30. Juvodden HT, Alnæs D, Agartz I, Andreassen OA, Server A, Thorsby PM, et al. Cortical thickness and sub-cortical volumes in post-H1N1 narcolepsy type 1: A brain-wide MRI case-control study. Sleep Med. 2024;116(February):81–9.

31. Abadi M, Barham P, Chen J, Chen Z, Davis A, Dean J, et al. TensorFlow: A System for Large-Scale Machine Learning.

32. Abadi M, Agarwal A, Barham P, Brevdo E, Chen Z, Citro C, et al. TensorFlow: Large-Scale Machine Learning on Heterogeneous Distributed Systems.

33. Keenan S, Hirshkowitz M. Sleep Stage Scoring. In: Principles and Practice of Sleep Medicine. Sixth Edit. Elsevier Inc.; 2017. p. 1567–75.

34. Trotti LM, Staab BA, Rye DB. Test-Retest Reliability of the Multiple Sleep Latency Test in Narcolepsy without Cataplexy and Idiopathic Hypersomnia. J Clin Sleep Med. 2013;9(8):789–95.

35. Lopez R, Doukkali A, Barateau L, Evangelista E, Chenini S, Jaussent I, et al. Test–Retest Reliability of the Multiple Sleep Latency Test in Central Disorders of Hypersomnolence. Sleep. 2017 Dec 1;40(12).

36. Roth T, Dauvilliers Y, Mignot E, Montplaisir J, Paul J, Swick T, et al. Disrupted nighttime sleep in narcolepsy. J Clin Sleep Med. 2013;9(9):955–65.

37. Fiorillo L, Monachino G, van der Meer J, Pesce M, Warncke JD, Schmidt MH, et al. U-Sleep’s resilience to AASM guidelines. Npj Digit Med. 2023;6:1–9.

38. Cesari M, Stefani A, Penzel T, Ibrahim A, Hackner H, Heidbreder A, et al. Interrater sleep stage scoring reliability between manual scoring from two European sleep centers and automatic scoring performed by the artificial intelligence-based Stanford-STAGES algorithm. J Clin Sleep Med. 2021;17(6):1237–47.

39. Fiorillo L, Pedroncelli D, Agostini V. Multi-scored sleep databases: how to exploit the multiple-labels in automated sleep scoring. 2023;(February):1–12.

40. Guillot A, Sauvet F, During EH, Thorey V. Dreem Open Datasets: Multi-Scored Sleep Datasets to Compare Human and Automated Sleep Staging. IEEE Trans Neural Syst Rehabil Eng. 2020;28(9):1955–65.

41. Moeller AL, Perslev M, Paulsrud C, Thorsen SU, Leonthin H, Debes NM, et al. Artificial intelligence or sleep experts: comparing polysomnographic sleep staging in children and adolescents. SLEEP. 2025 Feb 28;1–9.

42. Lee YJ, Lee JY, Cho JH, Choi JH. Interrater reliability of sleep stage scoring: a metaanalysis. J Clin Sleep Med. 2022;18(1):193–202.

43. Younes M, Hanly PJ. Minimizing Interrater Variability in Staging Sleep by Use of Computer-Derived Features. J Clin Sleep Med. 2016;12(10):1347–56.

44. Rosenberg RS, Van Hout S. The American Academy of Sleep Medicine inter-scorer reliability program: sleep stage scoring. J Clin Sleep Med. 2013;9(1):81–7.

45. Danker-Hopfe H, Kunz D, Gruber G, Klösch G, Lorenzo JL, Himanen SL, et al. Interrater reliability between scorers from eight European sleep laboratories in subjects with different sleep disorders. J Sleep Res. 2004;13(1):63–9.

46. Fiorillo L, Puiatti A, Papandrea M, Ratti PL, Favaro P, Roth C, et al. Automated sleep scoring: A review of the latest approaches. Sleep Med Rev. 2019;48:1–12.

