## Supplementary Material for "Optimizing automated sleep stage scoring of 5-second mini-epochs: a transfer learning study"

### General Scoring Rules

1. **Scrolling**: It is allowed to scroll back and forth within a 20-minute segment.
2. **Independent Scoring**: Each 5-second mini-epoch is scored as independently as possible; however, it is allowed to rely on context from previous or subsequent mini-epochs.
3. **Characteristics**: If one characteristic from a stage (listed below) is present in a given mini-epoch, that stage should be scored.
4. **Majority Rule**: If two or more stages are present in the same mini-epoch, score the stage that fills out the majority of the mini-epoch.
5. **No characteristics**: If no characteristics are present, there is the opportunity to mark this, however the context of surrounding mini-epoch along with pattern recognition should be used to decide which sleep stage it most certain is. This opportunity does not apply to wake and N3.
6. **Tonus**: Low tonus is not a specific characteristic as tonus can be low throughout all night. If tonus is used for pattern recognition, the context of the surrounding mini-epoch should be used to decide sleep stage.

### Stage characteristics

**Wake (W)**

- ≥50% Alpha activity.

**N1:**

- Slow eye movements.
- Vertex sharp waves.
- Note: If no slow eye movements or vertex sharp waves are seen in a given mini-epoch and no other characteristics suggesting other sleep stages are present and surrounding mini-epochs are scored as N1 and/or the background EEG resembles N1 or does not suggest another stage score “N1 without characteristics”.

**N2:**

- Sleep spindles.
- K-complexes.
- Note: If no sleep spindle or K-complex are seen in a given mini-epoch, and no other characteristics suggesting other sleep stages are present and surrounding mini-epochs are scored as N2 and/or the background EEG resembles N2 or does not suggest another stage score “N2 without characteristics”.

**N3:**

- High-amplitude, low-frequency delta waves (SWS).

**REM:**

- Rapid eye movements.
- Note: A rapid eye movement must be present in the mini-epoch for “REM” to be scored, otherwise if no other characteristics suggesting other sleep stages are present, and the surrounding mini-epochs are scored as REM, and/or the background EEG resembles REM or does not suggest another stage; score “REM without characteristics”.
- Note: If there is more than 50% alpha activity in the mini-epoch but no eye movements, however, it looks more like REM sleep than wake; “REM with alpha” should be scored.
- Note: If there is more than 50% alpha activity in the mini-epoch but eye movements and it looks more like REM sleep than wake; “REM” should be scored.

### Protocol

**Step 1:** The human scorer has 10 minutes to investigate a given full PSG in DOMINO.

**Step 2:** The human scorer scores mini-epochs of the first 20-minute segment from a given PSG in MATLAB following the rules above.

**Step 3:** The human scorer scores mini-epochs of the second 20-minute segment from a given PSG in MATLAB following the rules above.

The two segments from each PSG are in random order, meaning the first segment may originate from a later time in the night than the second segment. These three steps are repeated for all PSGs.

| Dataset | Total number of participants (n=100) | NT1 patients  (n=38) | Siblings  (n=62) |
| --- | --- | --- | --- |
| Training | 80 | 30 | 50 |
| Validation | 10 | 4 | 6 |
| Testing | 10 | 4 | 6 |

Table S1: Numbers of patients and siblings in the training, validation and test sets.
